# The Contribution of Parietal Cortex to Visual Salience

**DOI:** 10.1101/619643

**Authors:** Xiaomo Chen, Marc Zirnsak, Gabriel M. Vega, Eshan Govil, Stephen G. Lomber, Tirin Moore

**Affiliations:** Department of Neurobiology and Howard Hughes Medical Institute, Stanford University School of Medicine, Stanford, CA 94305, USA; Department of Physiology and Pharmacology, Department of Psychology, and Brain and Mind Institute, The University of Western Ontario, London, ON N6A 5K8, Canada

**Author notes:** Equal contribution (Xiaomo Chen, Marc Zirnsak). **Lead Contact:** Tirin Moore, Dept. of Neurobiology, Fairchild Bldg., 299 Campus Drive, Stanford University School of Medicine, Stanford CA 94305.

## Abstract

Unique stimuli stand out. In spite of an abundance of competing sensory stimuli, the detection of the most salient ones occurs without effort, and that detection contributes to the guidance of adaptive behavior. Neurons sensitive to the salience of visual stimuli are widespread throughout the primate visual system and are thought to shape the selection of visual targets. However, mechanisms underlying the representation of salience remain elusive. Among the possible candidates are areas within posterior parietal cortex, which appear to be crucial in the control of visual attention and are thought to play a unique role in representing stimulus salience. Here we show that reversible inactivation of parietal cortex not only selectively reduces the representation of visual salience within the brain, but it also diminishes the influence of salience on visually guided behavior. These results demonstrate a distinct contribution of parietal areas to vision and visual attention.

## Introduction

Throughout the brain, sensory input is continually filtered according to its relevance to behavioral goals, or according to its featural attributes and physical salience. Much progress has been made in identifying the neural circuits controlling goal-driven, or top-down, attention ^1^. In contrast, the mechanisms controlling salience-driven, or bottom-up, attention remain largely unknown. In the primate brain, the control of visual attention appears to be accomplished by neurons distributed within areas of prefrontal ^2–5^ and posterior parietal cortex (PPC) ^3, 4^ along with the superior colliculus ^6, 7^, and the pulvinar ^8, 9^. A lingering major question however is whether any of these structures contributes distinctively to bottom-up attention. Although many studies have examined the influence of visual salience on the responses of neurons in these structures ^3, 10–12^ and within posterior visual cortex ^13–18^, none have identified the structures necessary for the representation of visual salience. The timing of emergent visual salience signals within PPC suggest that neurons there may be causally involved ^3^, but this has yet to be explored. Here we tested the contribution of PPC to visual salience by reversibly inactivating it in behaving monkeys and measuring its effects both on the representation of salience among neurons with input from PPC, and on visually guided behavior.

## Results

### Behavioral effects of PPC inactivation

We reversibly inactivated large portions of PPC of two behaving monkeys (J and Q) via cryoloops which were chronically implanted within the intraparietal sulcus (IPS) (Methods)(Extended Data Figure 1). Cryoloops have been used extensively in the primate brain to temporarily eliminate the spiking activity of neurons within large expanses of neocortex in behaving animals ^19–22^. To assess the effectiveness of the inactivation, we first measured its impact on behaviors known to be affected by disruption of PPC activity in primates ^23, 24^. We did this in two ways. First, we measured the effects of inactivation on exploratory eye movements during free-viewing of complex images. Monkeys were allowed to freely view large images (79-98 by 49-55 degrees of visual angle, dva) for 3 seconds (Figure 1a). Consistent with the effects of parietal damage in human patients, inactivation of PPC in monkeys reduced the tendency to visually explore the contralateral half of space (Figure 1b and Extended Data Figure 2). To quantify this effect, we computed the density of fixations during free-viewing across all images for the two monkeys, and then compared the densities between control and inactivation (Figure 1c). For both monkeys, PPC inactivation reduced the fixation density within the contralateral visual field, resulting in a significant reduction in the proportion of fixations contralateral to the inactivation (monkey J, control_contra_= 0.49, inactivation_contra_= 0.37, *P*<10^−3^; monkey Q, control_contra_= 0.71; inactivation_contra_= 0.53, *P* <10^−34^) and a shift in the center of mass of fixations toward the ipsilateral visual field (monkey J, shift = 5.11 dva, *P*<10^−28^; monkey Q, shift = 4.14 dva, *P*<10^−28^). Thus, even with a coarse measure of behavior, the effect of PPC inactivation was clear.

**Figure 1.**
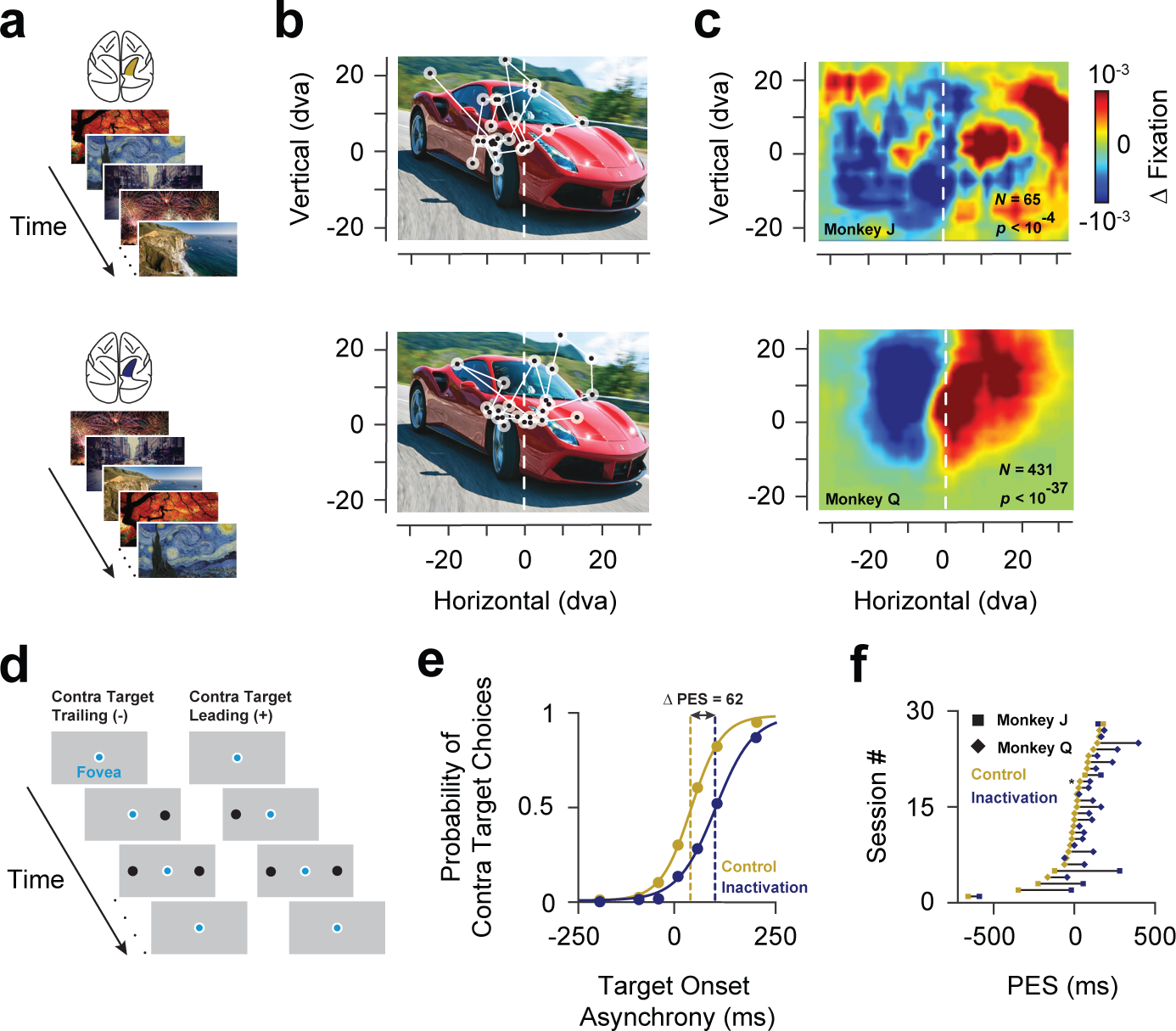
Behavioral effects of PPC inactivation. **a**. Free-viewing task. Images presented to the monkeys included real-word photographs, paintings, cartoons, and abstract patterns. Identical images were presented during both control (top, gold shading in IPS) and inactivation blocks (bottom, blue shading in IPS). Gold and blue shading in the brain icon denotes control and unilateral PPC inactivation, respectively. **b**. Example image presented to one monkey during a control (top) and inactivation block (bottom). Circles indicate regions of fixation and the lines indicate saccades. **c**. Change in fixation densities across the population of images for Monkey J (top) and Monkey Q (bottom). **d**. Double-target, choice task. Two targets were presented at varying temporal onset asynchronies within the ipsilateral and contralateral hemifields. **e.** Example experimental session for one monkey. Target choice functions during control and during PPC inactivation are plotted in gold and blue, respectively. **f**. Distribution of shifts in the PES across all sessions in the two monkeys.

**Figure 2.**
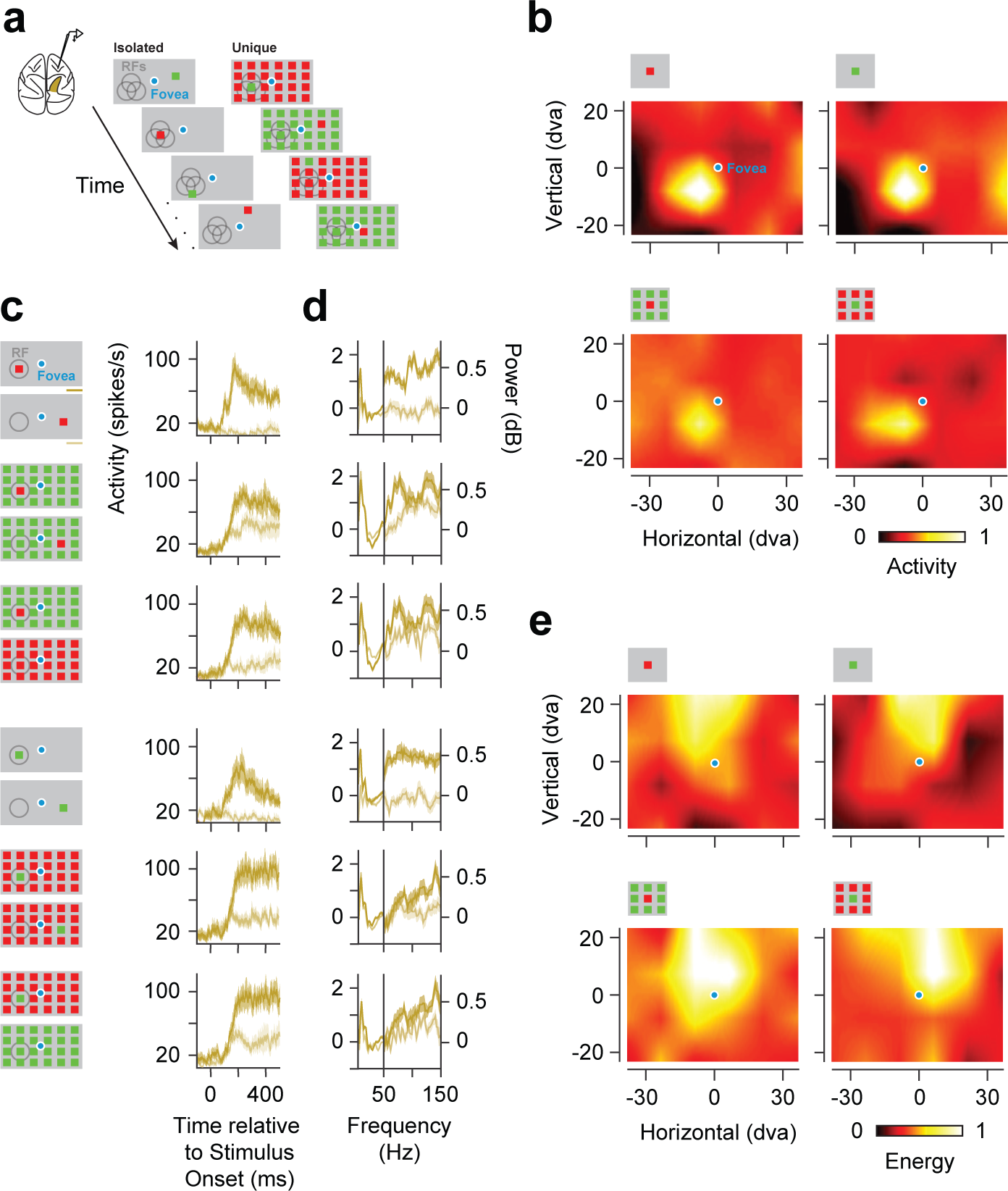
Prefrontal representation of visual salience in neuronal and LFP activity. **a**. Visual stimuli consisted of a single colored stimulus presented in isolation (Isolated), or among an array (6 × 4) of identically colored stimuli (Unique). **b**. Example CRFs of a single FEF neuronal recording mapped with an isolated red or green stimulus (top) and URFs of the same neuronal recording mapped with a unique red or green stimulus (bottom). **c.** Histogram of example neuronal responses to isolated and unique stimuli presented inside the CRF/URF (dark gold), shown with responses to single and unique stimuli presented outside of the CRF/URF or to identically colored stimulus arrays (light gold). **d**. Response spectra of an example FEF LFP recording. Same conventions as in c. **e**. High-gamma band CRFs and URFs for an example recording.

Second, we used a double-target, choice task to psychophysically assess the effect of inactivation on the tendency of monkeys to choose targets in the two hemifields ^25, 26^. In this task, monkeys were rewarded for choosing between two saccadic targets, one located within the contralateral hemifield, and one in the ipsilateral hemifield. The temporal onset of the two targets was systematically varied such that the contralateral stimulus could appear earlier or later than the opposite stimulus (Figure 1d). The monkey’s tendency to select the contralateral target could then be measured as the temporal onset asynchrony required for equal probability of selecting either target. Thus, a neglect of one hemifield would result in a shift of the point of equal selection (PES) toward the ipsilateral hemifield. Indeed, that is what we observed; the PES shifted in favor of the ipsilateral target (Figure 1e). As a result, in order for contralateral targets to be chosen as frequently, they needed to appear earlier than during control blocks. This effect was reliably obtained in both monkeys (monkey J, Δ_PES_=189.19 ± 76.13 ms, *P* < 0.04; monkey Q, Δ_PES_= 85.19 ± 13.56 ms, *P* < 2.26 × 10^−6^) (Figure 1f). Notably, inactivation of the ventral IPS alone was sufficient to produce effects equivalent to both dorsal and ventral inactivation (Extended Data Figure 3), consistent with an earlier comparison of dorsal and ventral lateral intraparietal area (LIP) ^27^. Overall, the magnitude of the observed effects was larger than that of previous studies using more localized PPC inactivations ^24, 25^. Thus, by both behavioral measures, PPC inactivation produced robust effects generally resembling the effects of PPC damage in monkeys ^23^, and were similar to hemispatial neglect in humans ^28^.

**Figure 3.**
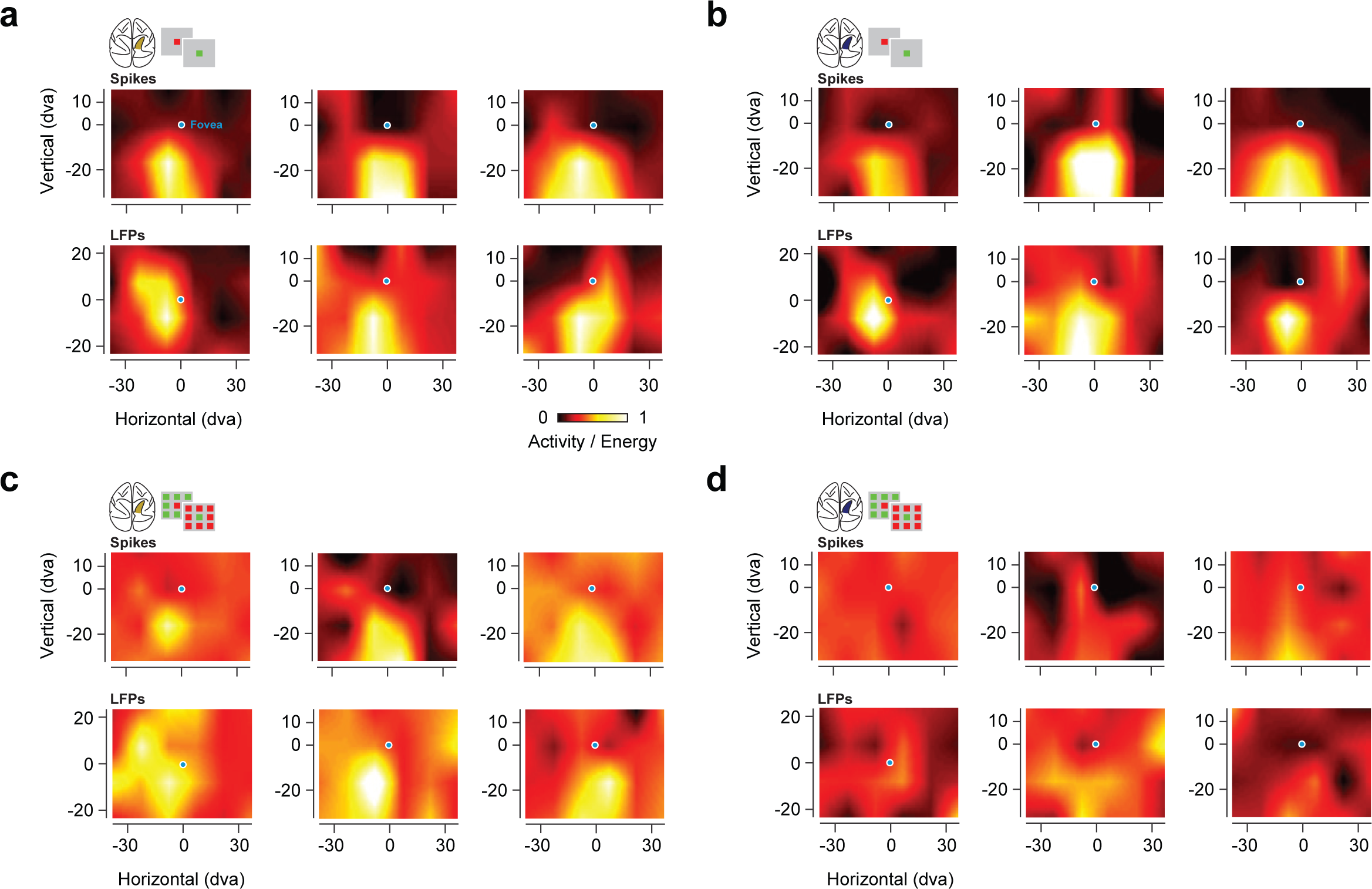
Prefrontal CRFs and URFs during PPC inactivation. **a**. Six example CRFs derived from spiking (top 3) and from high-gamma band LFPs (bottom 3) during control. **b.** Same RFs as in **a**, but during PPC inactivation. **c.** Six example URFs derived from spiking (top) and from high-gamma band LFPs (bottom) during control. **d.** Same RFs as in **c,** but during PPC inactivation.

### Representation of salience by neurons within prefrontal cortex

To assess the effects of PPC inactivation on the representation of visual salience, we recorded neuronal spiking activity and local field potentials (LFPs) within prefrontal cortex, specifically within the frontal eye field (FEF). Neurons within the FEF receive input directly from most areas within posterior visual cortex ^29^, as well as strong inputs from areas within PPC, particularly LIP ^30^. We recorded the activity of FEF neurons in two behaving monkeys using multichannel microelectrodes (Methods). We then assessed the representation of visual salience in the recorded neuronal (n=352) and LFP activity (n=192) that was present prior to inactivation. For both types of activity, we measured the responses to visual stimuli consisting of a single colored stimulus presented in isolation, or among an array of identically or differently colored stimuli (Figure 2a).

Using an isolated red or green stimulus, we mapped the region of space most sensitive to visual stimulation, i.e., the classical receptive field (CRF), for each neuronal recording. FEF neurons are not typically selective for stimulus features, including color ^31, 32^, as in the example shown in Figure 2b. Nonetheless, FEF neurons are sensitive to stimuli that are unique among competing ones ^3, 12^. Thus, for each neuronal recording, we could also map the region of space most sensitive to a unique stimulus (URF). Neurons therefore signaled the location of both isolated and unique stimuli, independent of color (figure 2b). Across our population of neurons, the difference in responses to an isolated red or green stimulus was typically small (median = 4.6%), consistent with previous studies ^31^. Nonetheless, for the same population, neuronal responses were robustly enhanced by the appearance of a unique stimulus in the URF. The enhancement was evident in comparisons with responses to arrays in which the unique stimulus fell outside of the URF (Unique_In_ – Unique_Out_). The enhancement was also evident in comparisons with responses to an array that rendered the URF stimulus identical to surrounding stimuli (Unique_In_ – Identical)(figure 2c). We quantified the two types of enhancement by computing a standard index of response enhancement, specifically the difference between the Unique_In_ and the Unique_Out_ (or Identical) responses, divided by their sum. Across the population, both types of enhancement were highly significant (median Unique_In_ – Unique_Out_ index = 0.11, *P* < 10^−45^; median Unique_In_-Identical index = 0.11, *P* < 10^−44^), with more than half of the population exhibiting significant effects in both of the comparisons (Unique_In_ – Unique_Out,_ 193/352; Unique_In_-Identical, 173/352).

In addition, we probed the representation of visual salience in the FEF LFPs. Information about the location of isolated visual stimuli is robustly signaled within the alpha (8-12Hz) and high-gamma (60-150Hz) bands of FEF LFPs ^33^. In the present study, we observed that activity in the high-gamma band, but not the alpha band, also robustly signaled the location of a unique visual stimulus (Extended Data Table 1). Compared to other frequency bands, responses to unique stimuli were most consistent in the high-gamma LFPs, and they were enhanced relative to responses to the appearance of unique stimuli outside of the URF and to arrays that rendered the URF stimulus identical to surrounding stimuli (Figure 2d). Using the high-gamma signal, we could derive visual RFs for both the isolated and the unique stimulus, similar to the spiking activity (Figure 2e). Across the population of recorded high-gamma LFPs, we observed both types of enhancement that we observed in the spiking responses (Unique_In_ – Unique_Out,_ median DEnergy = 0.50 dB, *P* < 10^−20^; Unique_In_ – Identical, median DEnergy = 0.47 dB, *P* < 10^−21^). Thus, similar to the spiking activity, the high-gamma LFPs were highly sensitive to visual salience.

### Salience signals in prefrontal cortex during PPC inactivation

Given the clear behavioral effects we observed during PPC inactivation, we next asked whether removing parietal input alters visual responses in the FEF. We reasoned that if indeed parietal areas contribute distinctively to the representation of visual salience, then PPC inactivation should selectively reduce salience signals downstream in the FEF. Indeed, that is what we observed. First, PPC inactivation did not change the selectivity of FEF neurons to color (paired t-test, *P* = 0.99). Second, it had only minimal effects on CRFs derived from spiking or LFP activity. However, inactivation dramatically altered URFs; that is, it diminished receptive fields mapped with a unique stimulus (Figure 3). During inactivation, visually driven activity was generally reduced in proportion to the magnitude of visual responses during control trials (ANCOVA main effect, *P* < 10^−41^). However, the size of the reduction significantly depended on the stimulus condition (ANCOVA interaction, *P* < 0.002)(Extended Data Table 2), with the URF stimulus yielding the greatest reduction in visual responses. This selective reduction can be seen in the example neuron shown in Figure 4a. In this example, responses to an isolated stimulus were virtually unaffected by PPC inactivation. In contrast, responses to the unique stimulus were robustly diminished compared to responses to arrays in which the unique stimulus fell outside of the URF, or an array that rendered the URF stimulus identical to surrounding stimuli. As a consequence of the selective reduction in visual responses, the two types of salience enhancement observed in FEF neurons were markedly reduced by inactivation (Control: Unique_In_ – Unique_Out_ index = 0.26, Inactivation: Unique_In_ – Unique_Out_ index = 0.10; Control: Unique_In_ – Identical index = 0.33, Inactivation: Unique_In_ – Identical index = 0.17). This pattern of results was similar across the population. For modulated neurons (n=193), inactivation reduced both types of enhancement by ∼38% (Control: median Unique_In_ – Unique_Out_ index = 0.21, Inactivation: median Unique_In_ – Unique_Out_ index = 0.13, *P* < 10^−9^; Control: Unique_In_-Identical index = 0.18, Inactivation: Unique_In_-Identical index = 0.11, *P* < 10^−13^).

**Figure 4.**
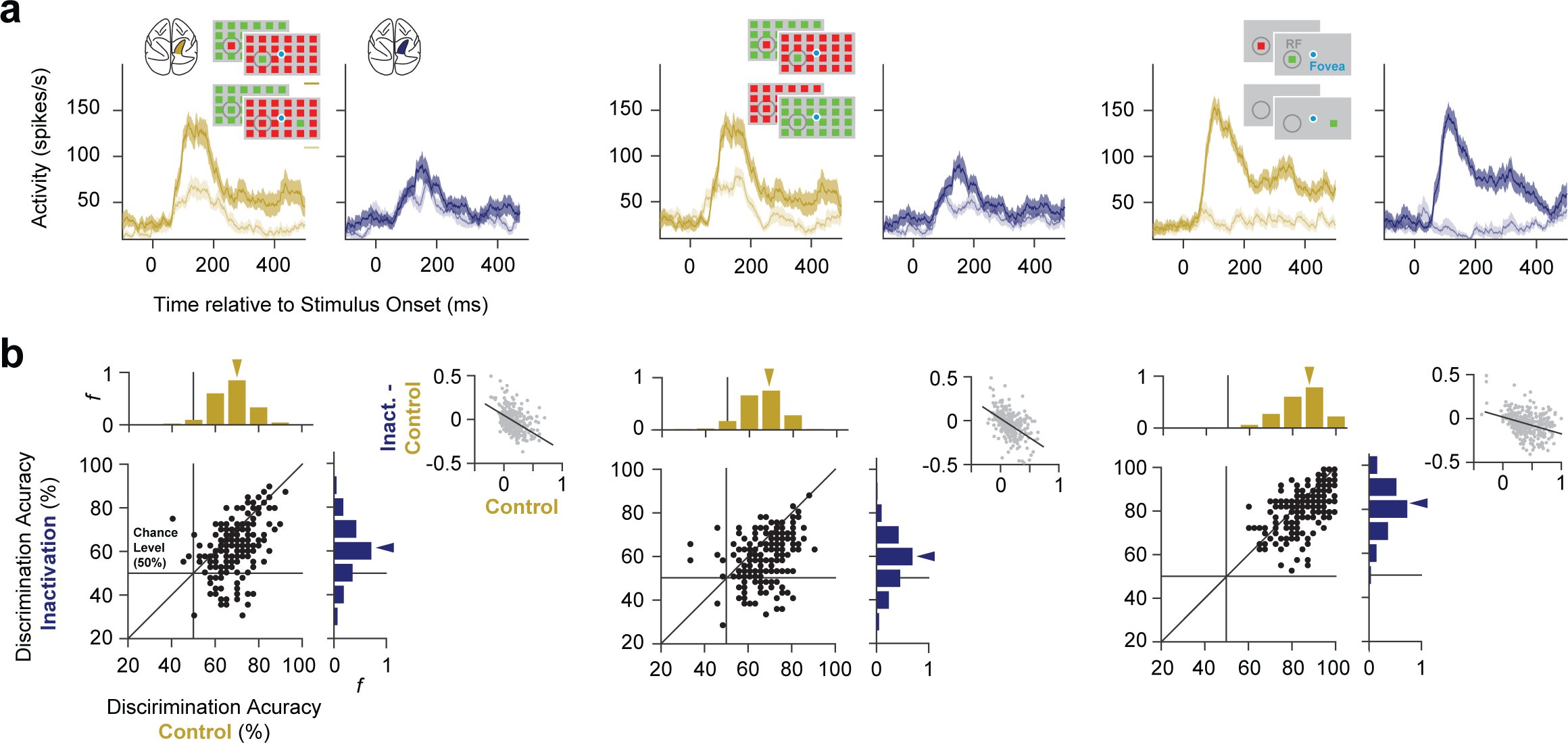
Representation of salience in prefrontal neuronal activity during PPC inactivation. **a**. Mean responses of an example neuron to different stimulus conditions during control (gold) and PPC inactivation (blue). Left, responses to Unique_In_ (dark) and Unique_Out_ (light) stimuli. Middle, responses of the same neuron to Unique_In_ (dark) and Identical (light). Right, responses of the same neuron to the isolated stimuli presented inside (dark) and outside of the CRF (light). **b**. Accuracy of classifiers trained on neuronal spiking activity to discriminate between different stimulus conditions during control and PPC inactivation. Left, accuracy of classifiers trained to discriminate between Unique_In_ and Unique_Out_ stimuli. Middle, accuracy of classifiers trained to discriminate between Unique_In_ and Identical stimuli. Right, accuracy of classifiers trained to discriminate between Isolated stimuli appearing inside and outside of the CRF. Scatter plots and marginal distributions compare discrimination accuracies across stimulus selective neuronal recordings (n = 193) during control and inactivation. Gray scatterplots show the reduction in enhancement indices during inactivation as a function of enhancement indices measured during control for all recordings (n = 352). Black lines show the linear regression fits.

To quantify the effects of PPC inactivation across the population of FEF neurons, we measured the accuracy of a linear classifier in discriminating between visual stimulus conditions using the trial-by-trial responses of each individual neuronal recording. We focused our analysis on 193 FEF neuronal recordings with significant response differences between the inside and outside RF conditions for both isolated and unique stimuli (see Methods) (Figure 4). For these neurons, sensitivity to visual salience was selectively reduced. During control trials, the classifier performed above chance in discriminating the unique inside and outside conditions (Unique_In_ vs. Unique_Out_) in 147 neuronal recordings. However, during inactivation that number was reduced by 40% to 88 (McNemar’s chi-square = 43.7, *P* < 10^−10^), and the median classifier performance was reduced by 7.5%, a 39% reduction in above-chance performance (see Methods). Similarly, the classifier performed above chance in discriminating between the unique RF stimulus and the identical array (Unique_In_ vs. Identical) in 135 neuronal recordings during control trials. Yet during inactivation that number was reduced by 44% to 75 (McNemar’s chi-square = 37.8, *P* < 10^−9^), and the median classifier performance was reduced by 7.5%, a 43% reduction in above-chance performance. This reduction in discrimination performance was accompanied by a reduction in the two types of salience enhancement, a reduction that was correlated with enhancement during control trials (Unique_In_ – Unique_Out_:, r = −0.40, *P* < 10^−17^; Unique_In_ – Identical: r = −0.44, *P* < 10^−20^). The slopes of both correlations were significantly steeper than that observed for responses to isolated stimuli (Unique_In_ – Unique_Out_ vs. Isolated: Dslope = −0.21, *P* < 10^−3^; Unique_In_ – Identical vs. Isolated: Dslope = −0.23, *P* < 10^−5^), which again indicates that the reduction in selectivity was larger for unique stimuli. Correspondingly, the reduction in performance during inactivation for classifiers trained to discriminate an isolated stimulus inside versus outside of the CRF was 2.9%, an 8% reduction in above-chance performance, which was significantly smaller than the reduction observed for unique stimuli (Unique_In_ – Unique_Out_ vs. Isolated: *P* < 10^−4^; Unique_In_ – Identical vs. Isolated: *P* < 10^−4^). Thus, PPC inactivation selectively reduced the representation of visual salience by neurons in prefrontal cortex.

Next, we examined the effects of PPC inactivation on the salience-driven enhancement of FEF LFP activity. During control trials, the enhancement in high-gamma band LFP responses to unique stimuli emerged ∼100 ms after the visual onset response and was evident in both Unique_In_ – Unique_Out_ and Unique_In_ – Identical comparisons (Figure 5). During PPC inactivation, we found that both types of enhancement were largely eliminated. As with the spiking activity, the reduction in high-gamma band responses to visual stimulation was largest when the unique stimulus appeared in the URF (Extended Data Figure 4). Consequently, both types of enhancement observed in high-gamma responses to unique stimuli were reduced during PPC inactivation (Unique_In_ – Unique_Out_: 37%, *P* < 0.003; Unique_In_ – Identical: 35%, *P* < 0.004). As with the spiking activity, smaller changes were observed in responses to isolated stimuli. Unlike responses to unique RF stimuli, in which only the high-gamma band responses discriminated between the inside and outside RF conditions, both the alpha and high-gamma band responses discriminated between the two for isolated stimuli. During inactivation, both signals remained. But more importantly, the difference between inside and outside RF responses in the high-gamma band were reduced to a lesser extent than that observed for unique RF stimuli (15%, *P* < 0.006; Unique_In_ – Unique_Out_ vs. Isolated: *P* < 0.02; Unique_In_ – Identical vs. Isolated: *P* < 0.03). So, as with the spiking responses, PPC inactivation selectively reduced the representation of visual salience in prefrontal LFPs.

**Figure 5.**
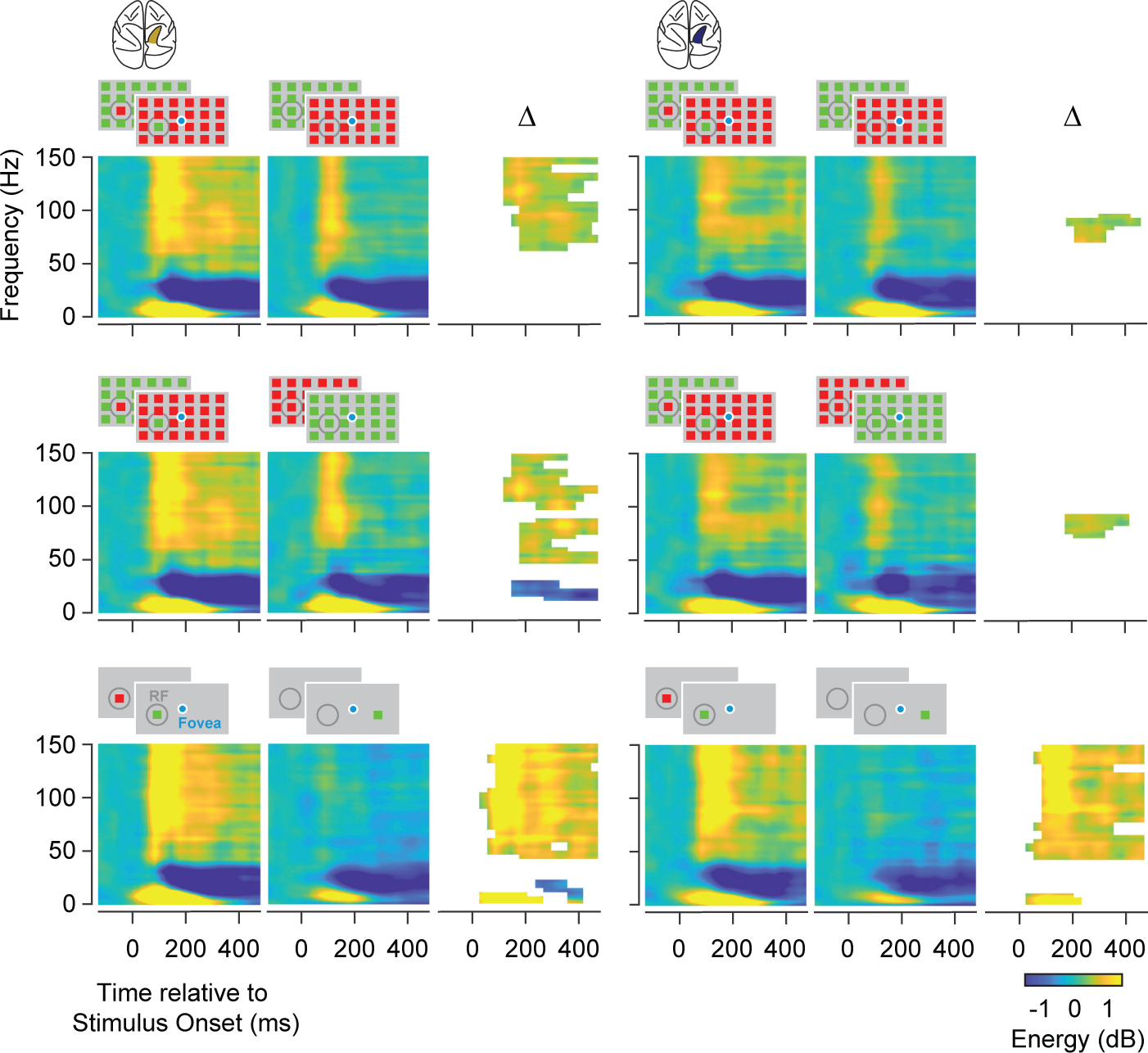
Representation of salience in prefrontal LFPs during PPC inactivation. Comparison of LFP time-frequency power spectrograms during control (left) and PPC inactivation (right). The first row compares the spectrograms of responses to Unique_In_ and Unique_Out_ stimuli, and their differences (Δ) during control and inactivation. The second row compares responses to Unique_In_ and Identical stimuli, and the third row compares responses to Identical stimuli presented inside or outside of the CRF. Difference plots only show time-frequency bins with significant energy differences.

### Changes in Salience-driven behavior during PPC inactivation

Given the selective reduction in the sensitivity of FEF neurons to visual salience during PPC inactivation, we wondered if there might be corresponding changes in salience-driven behavior. Since the FEF has a well-established role in the programming and triggering of visually guided saccadic eye movements ^34, 35^, we considered that the inactivation might alter the influence of salience on this behavior. Our initial behavioral results with the free-viewing and double-target tasks indeed revealed robust effects of PPC inactivation on visually guided eye movements. However, the free-viewing task provided an additional opportunity to assess whether the inactivation altered the influence of visual salience on eye movements. Beginning with the earliest model ^36^, a wealth of models have been developed to quantify physical salience within images based on the contrast across various feature dimensions (e.g. color) ^37–40^, thereby identifying points of relative salience within an image. Moreover, these models can be used to predict where in the image human observers fixate with varying accuracy ^40^. We leveraged this approach to quantify the distribution of salience within the images our monkeys freely viewed, and to assess the influence of salience on eye movements. Salience ‘maps’ were computed from each of the 487 images viewed by the two monkeys (65, monkey J; 431, monkey Q) using the Graph-based Visual Salience model (GBVS) ^37^ (Figure 6a). Next, as in human studies, we measured the 2D correlation between the distribution of fixations and the salience map of each image, before and after PPC inactivation (Methods)(Figure 6b). Prior to inactivation, as in human observers, fixations were weakly, but significantly, correlated with image salience _40_ (Monkey J, r_median_= 0.11, *P* <10^−28^; Monkey Q, r_median_= 0.15, *P* < 10^−172^). Moreover, for both monkeys, PPC inactivation significantly reduced the correlations for fixations made throughout the freely viewed images (Monkey J, Δr_median_ = −0.03, *P* < 10^−4^; Monkey Q, Δr_median_ = −0.01, *P* < 10^−5^), indicating that inactivation diminished the influence of salience on visually guided eye movements. More importantly, the reduced correlations with salience were observed within the contralateral space in both monkeys. We examined the change in correlations separately for ipsilateral and contralateral fixations, defined either in eye-centered or in head-centered coordinates (Figure 6b). In the eye-centered analysis, we divided fixations within each image into those resulting from movements made in a direction contralateral or ipsilateral to the PPC inactivation, and we computed 2D correlations separately for the two sets of fixations. This analysis revealed that PPC inactivation reduced correlations for contralaterally directed fixations in both monkeys (Monkey J, Δr_median_ = −0.02, *P* < 0.008; Monkey Q, Δr_median_ = −0.02, *P* < 10^−13^) (Figure 6c). In the head-centered analysis, we divided fixations within each image into those that landed within the contralateral or ipsilateral side of the image, regardless of the movement direction (Figure 6b). Similar to the eye-centered results, the image-centered analysis revealed that PPC inactivation reduced correlations for fixations within the contralateral half of images in both monkeys (Monkey J, Δr_median_ = −0.02, *P* < 0.002; Monkey Q, Δr_median_ = −0.03, *P* < 10^−17^) (Figure 6c). By comparison, we observed no consistent changes within the ipsilateral hemifield (Extended Data Figure 5). The pattern of results was similar when image salience was computed with another popular model^38^. Importantly, the consistent decrease in contralateral correlation coefficients we observed was not a result of decreased saccadic accuracy during inactivation, as we did not observe such an effect (Extended Data Figure 6). Instead, the decreased correlations appeared to result from a reduced influence of visual salience on the pattern of fixations directed toward the contralateral visual space, and fixations made within the contralateral half of images during PPC inactivation, consistent with the neurophysiological results.

**Figure 6.**
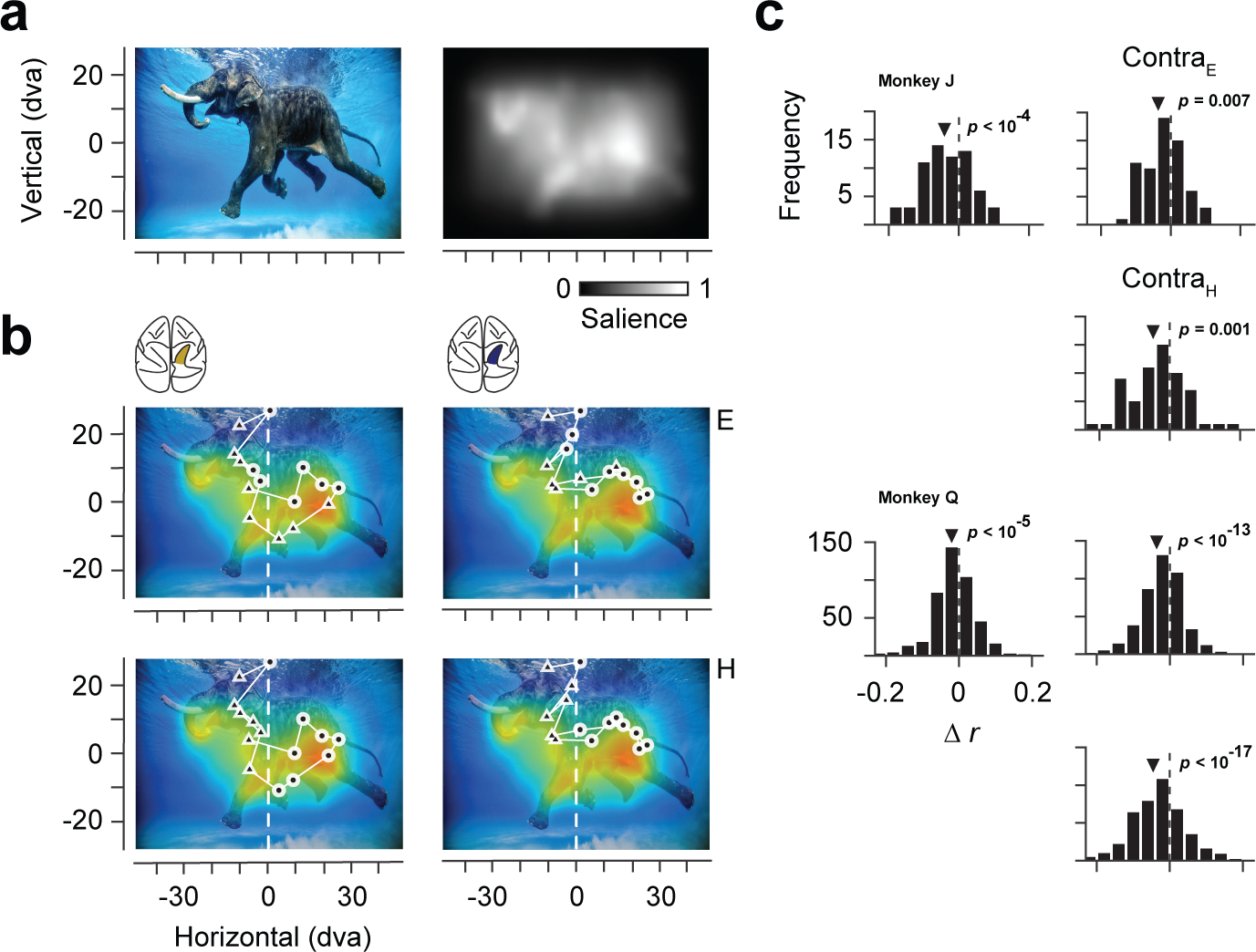
Changes in salience-guided fixations during PPC inactivation. **a**. Example image from the free viewing task (top left) and corresponding salience map (top right). **b**. Correspondence between salience and fixations made in the example image before and after PPC inactivation. Top row fixations are labelled in eye-centered coordinates as contralaterally (triangles) or ipsilaterally (circles) directed movements. Bottom row shows the same fixations labelled in head-centered coordinates as landing in the contralateral or ipsilateral half of the image. **c**. Distribution of changes in fixation-salience map correlation coefficients (*r*_inactivation_ – *r*_control_) across the population of images for the two monkeys. Left histograms show distributions based on coefficients measured from fixations across the full image. Right histograms show distributions based on contralateral fixations, defined in eye-centered (Contra_E_) or head-centered (Contra_H_) coordinates.

## Discussion

A wealth of neurophysiological studies have provided strong evidence of a distinct role of PPC in the representation of visual salience ^3,41^. However, a causal test of that role has been lacking. Our results indicate that the representation of salience within PPC is necessary for the emergence of salience signals in prefrontal cortex and for the influence of salience on behavior. During PPC inactivation, we observed that neural responses to unique visual stimuli were reduced relative to responses to non-unique or isolated stimuli. Furthermore, we found that these reductions in neural signals were accompanied by impairments in salience-guided behavior. Parietal cortex, which is extensively evolved and enlarged in primates, consists of a constellation of multimodal, integrative cortical areas involved in the transformation of sensory and motor signals across different coordinate frames and motor effectors ^42^. Within the visual domain, PPC areas such as area LIP are heavily interconnected with feature-selective areas within extrastriate visual cortex, where salience in each feature dimension is thought to be computed ^43^. Thus, areas like LIP might integrate salience across multiple features in order to select unique stimuli and guide bottom-up attention and behavior.

Although both the neurophysiological and behavioral impairments were robust, they were not absolute, as is often the case with studies using inactivation or lesions to probe mechanisms of visual perception ^21, 44–46^. Thus, it is important to consider which mechanisms or structures might underlie the residual function we observed. The FEF clearly depends on input from PPC, but the residual representation of salience there could be computed within the FEF, particularly given the FEF’s direct connections with feature-selective extrastriate visual areas ^29^. Alternatively, recent studies have identified representations of visual salience with very short latencies within the superficial, visual layers of the superior colliculus ^47^, which is heavily connected with the FEF, and is involved in the control of visually guided eye movements. Indeed, studies in birds reveal an important role of the midbrain in the representation of stimulus salience ^48^. In addition, biologically plausible models of the computation of visual salience highlight the necessary role of visual cortical areas in generating feature contrast ^36, 38, 43^, and therefore inactivation of PPC should not be expected to completely eliminate those signals in downstream areas like the FEF. Indeed, as with top-down visual attention, which is controlled by a set of distributed structures^1^, it appears unlikely that salience-driven, bottom-up attention is controlled by a single brain area. Nonetheless, our results identify areas within PPC as being necessary for that basic function.

## Acknowledgements

We thank E. Margalit and M. Plitt for assistance with parts of the behavioral analyses. This work was supported by NIH EY014924 to T.M.

## METHODS

### General and Surgical Procedures

Two male rhesus monkeys (Macaca mulatta, 17 and 16 kg), monkey J and monkey Q, were used in these experiments. All experimental procedures were in accordance with National Institutes of Health Guide for the Care and Use of Laboratory Animals, the Society for Neuroscience Guidelines and Policies, and Stanford University Animal Care and Use Committee. Surgery was conducted using aseptic techniques under general anesthesia (isoflurane) and analgesics were provided during postsurgical recovery. Each animal was surgically implanted with a titanium head post and a cylindrical titanium recording chamber (20 mm diameter) overlaying the arcuate sulcus. A craniotomy was then performed in the chambers on each animal, allowing access to the FEF.

### Cryoloops surgery and reversible inactivation of PPC

Each animal was surgically implanted with two stainless steel cryoloops within the intraparietal sulcus of one hemisphere. The size and shape of the cryoloops were customized to fit the contours of the IPS and to completely fill the sulcus. One longer loop (2.2-2.4×0.4 cm) was placed ventrally, and one shorter loop was placed dorsally (1.7-1.8×0.3 cm) (Extended Figure 1). During the cryoloop surgery, unilateral craniotomies were made over the intraparietal sulcus. Cryoloops were then placed beneath the dura and upon the surface of the arachnoid membrane in the dorsal and ventral intraparietal sulcus. The loops were secured to the skull with bone screws and dental acrylic. The dura was replaced and bone defects around the implanted cooling loops were repaired with original bone, Gelfoam (Pfizer) and dental acrylic. For detailed cryoloop implantation procedures, see Lomber and Payne^49^.

### Inactivation procedures

Cortex within the IPS was cooled by pumping chilled methanol through the loop tubing. Loop temperature was monitored and accurately regulated within 1 °C of the desired value by controlling the rate of methanol flow. A stable loop temperature (around 5 °C) was reached in ∼5-10 min of initiating cooling, and normal brain temperature was regained in ∼2 min after the cessation of cooling as a result of the infusion of warm blood ^50,51^. Loop temperatures around 5 °C reliably deactivate the full thickness of underlying cortex ^49^. During experimental sessions, blocks of cryo-inactivation lasted 30-60 minutes.

### BEHAVIOR

#### Behavioral tasks: free-viewing

During all behavioral measurements eye position was monitored and stored at 1000 Hz (Eyelink 1000, SR Research). While seated and head-restrained, monkeys were rewarded for freely viewing complex images, similar to a previous study ^52^. Images (Monkey J: 79×49 dva; Monkey Q: 98×55 dva;) were presented on a display (Monkey J: Samsung 2233RZ, 120 Hz refresh rate, 1680 × 1050 pixel resolution; Monkey Q: ASUS VS228; 75 Hz refresh rate, 1920 × 1080 pixel resolution) positioned 28-30 cm in front of the animal. A novel set of 100 images was used for each experimental session. In each trial, monkeys fixated a central fixation point (1×1 dva fixation window) on a gray background (60 cd/m^2^) for 500 ms to initiate the image presentation. Each image was displayed for 3 seconds and was shown in both control and inactivation blocks. Monkeys were rewarded at the end of each trial for exploring the image for the full presentation time. The sequence of control and inactivation blocks was varied across experimental sessions.

#### Behavioral tasks: choice task

To measure the effects of PPC inactivation on target selection, we quantified the monkey’s tendency to select stimuli at a particular location as the target of a saccadic eye movement. We employed a double-target, choice task similar to one used previously ^53^. In the task, the monkey was rewarded for making saccades to either one of two visual stimuli (1 dva diameter) appearing at diametrically opposed locations on the same display as used in the free-viewing task. One of the stimuli was positioned within the contralateral hemifield, and the other in the ipsilateral hemifield. The appearance of the two stimuli on a given trial occurred within a range of temporal onset asynchronies (TOAs), from trials in which the contralateral target appeared first (positive TOAs) to trials in which the contralateral target appeared second (negative TOAs). The range of TOAs for a given block of trials was −800 to 800 ms, with 7-9 discrete TOAs evenly spaced within that range, including zero. Trials were randomly interleaved such that on any given trial the monkey could not predict the TOA. In a given experimental session, at least 2 blocks of trials were collected, one prior to PPC inactivation, and one following it. Each block consisted of at least 10 trials per TOA. Each pair of pre- and post-inactivation target selection blocks could be used to compare the probability that the monkey would choose one target over the other as a function of TOA. We used logistic regression, on a trial-by-trial basis ^54^ to estimate the point of equal selection (PES) ^53^, that is the TOA for which the selection of either target has equal probability.

#### Salience map and correlation analysis of fixations during free-viewing

Both the graph based visual salience model (GBVS) ^37^ and the Itti-Koch-Niebur model ^38^ were used to compute the salience map for each image. For both models, feature channels including color, luminance, and orientation were used in the computation of the salience map.

Similar to human free viewing studies ^40^, a 2-D Pearson’s correlation coefficient was computed to quantify the relationship between the salience map *SM_i_* of image *i* and the fixation density map *FDM_i_* of image *i*. Raw salience maps (32 × 18, Δ*x* = {2.5, 3.0} dva, Δ*y* = {2.7, 3.0} dva, depending on the display) were used without interpolation. Fixation density maps were calculated in the same spatial resolution as the salience maps. The correlation for the full *SM* and full *FDM* is defined as

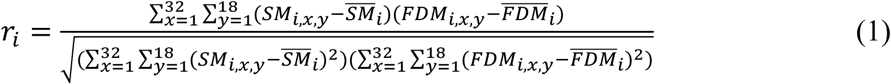

where *SM_i,x,y_* denotes the salience map for the *i*th image at the location (*x*, *y*), *FDM_i,x,y_* denotes the fixation density at the location (*x*, *y*) when the monkey was viewing the *i*th image, 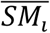 denotes the mean salience across the whole image, and 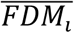 denotes the mean fixation density across the whole image. For correlations in eye-centered coordinates, the *FDM* was computed separately for all eye movements that had a contralateral and ipsilateral component. For correlations in head-centered coordinates, the *SM* and *FDM* were computed separately for contralateral and ipsilateral halves of each image. Results of comparisons of correlations across control and inactivation blocks yielded similar results when using all fixations or matched numbers of fixations between the two blocks.

### ELECTROPHYSIOLOGY

#### FEF recording procedures

Recording sites within the FEF were identified by eliciting short-latency, fixed vector saccadic eye movements with trains (50-100ms) of biphasic current pulses (≤50 µA; 250 Hz; 0.25 ms duration) as in previous studies (Bruce and Goldberg, 1985). Single-neuron and local field potential (LFP) recordings were obtained with 16 or 32-channel linear array electrodes with contacts spaced 150 µm apart (V and S-Probes, Plexon, Inc). Electrodes were lowered into the cortex using a hydraulic microdrive (Narishige International). Neural activity was measured against a local reference, a stainless guide tube, which was close to the electrode contacts. At the preamplifier stage, signals were processed with 0.5 Hz 1-pole high-pass and 8 kHz 4-pole low-pass anti-aliasing Bessel filters, and then divided into two streams for the recording of LFPs and spiking activity. The stream used for LFP recording was amplified (×500 – 2000), processed by a 4-pole 200 Hz low-pass Bessel filter and sampled at 1000 Hz. No other filters were used in the analyses. The stream used for spike detection was processed by a 4-pole Bessel high-pass filter (300 Hz) a 2-pole Bessel low-passed filter (6000 Hz), and was sampled at 40 kHz. Extracellular waveforms were classified as single neurons or multi-units using online-template-matching and subsequently confirmed using offline sorting (Plexon).

#### CRF and URF measurements

We measured LFP and spiking activity derived CRFs within the FEF by randomly presenting a single isolated probe stimulus out of a 6×4 probe grid extending 75×45 dva (Isolated stimulus condition). In each recording session, we placed the probe grid so as to cover the area where we expected to find most RF locations based on the saccade vectors evoked by electrical stimulation at a given recording site. The probes consisted of fully saturated red or green 7×7 dva squares. Similarly, we measured LFP and spiking activity derived URFs within the FEF by randomly presenting a uniquely colored probe stimulus among an array of differently colored stimuli, either a single green among 23 red or a single red among 23 green stimuli (Unique stimulus condition). In addition, we also measured neural response (LFP and spiking activity) to an identically colored (24 red or 24 green) stimulus array (Identical stimulus condition). Each stimulus condition was repeated at least 8 times during both control and inactivation conditions. Different stimulus conditions were pseudo-randomly interleaved.

In each trial, monkeys were required to fixate a central fixation point (1×1 dva fixation window) on a gray background (60 cd/m^2^) for 500 ms to initiate the trial. Subsequently, either an Isolated, Unique, or Identical stimulus was presented for 500 ms while the monkey maintained fixation. Following stimulus offset, the monkey received a juice reward after an additional 300ms of fixation.

#### Enhancement Index

We used two indices to quantify the enhancement of neuronal responses to Unique stimuli appearing inside the URF,

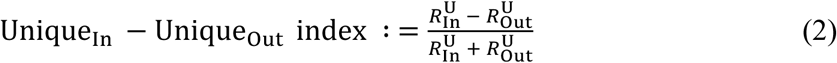

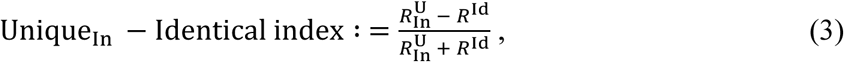

with 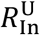 denoting the mean neuronal response to Unique stimuli presented inside the URF, [0, 500) ms relative to stimulus onset, 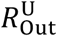 denoting the mean response to Unique stimuli presented outside the URF, and *R*^Id^ denoting the mean response to Identical stimulus arrays.

#### Support Vector Machine (SVM) linear classifier

We used a linear support vector machine (SVM) ^55^ to quantify the selectivity of neurons to Unique and Isolated stimuli. A classifier was trained to discriminate between neuronal responses to Unique stimuli presented inside the URF and responses to Unique stimuli presented outside the URF, and between responses to Identical stimuli, on a trial-by-trial basis. Similarly, a classifier was trained to discriminate neuronal responses to Isolated stimuli presented inside the CRF from responses to Isolated stimuli presented outside the CRF, on a trial-by-trial basis. Before training, spike counts for each neuronal recording were normalized across all stimulus conditions. All reported discrimination accuracies are based on four-fold cross-validation. Permutation tests (1000 repetitions) were used to determine whether the discrimination accuracy of a given neuronal recording was significantly greater than that expected by chance (discrimination performance of the classifier after label shuffling).

#### Time-frequency analysis

Matching pursuit (MP) decomposition was used in calculating the spectrogram to optimize temporal and frequency resolutions ^56, 57^. This multiscale decomposition allows sharp transients in the LFP signal to be represented by functions that have narrow temporal support, rather than oscillatory functions with a temporal support of hundreds of milliseconds. The algorithm is an iterative procedure that selects a set of Gabor functions (atoms) from a redundant dictionary of functions that constitute the best possible description of the original signal. Time–frequency plots were then obtained by calculating the Wigner distribution of every atom and taking the weighted sum. We performed the MP computation using custom MATLAB (MathWorks) scripts and the MP toolbox (https://github.com/supratimray/MP) ^56^. Permutation tests (N=1000) with multiple correction were used to determine whether the energy distribution at selected times and frequencies were significantly different between stimulus conditions. The mean LFP power for each frequency band (alpha, 8-12Hz; beta, 12-30Hz; low gamma, 30-60Hz; and high gamma, 60-150 Hz) was calculated as the mean of the energy [0, 500) ms after visual stimulus onset.

---

**Extended Data Figure 1.**
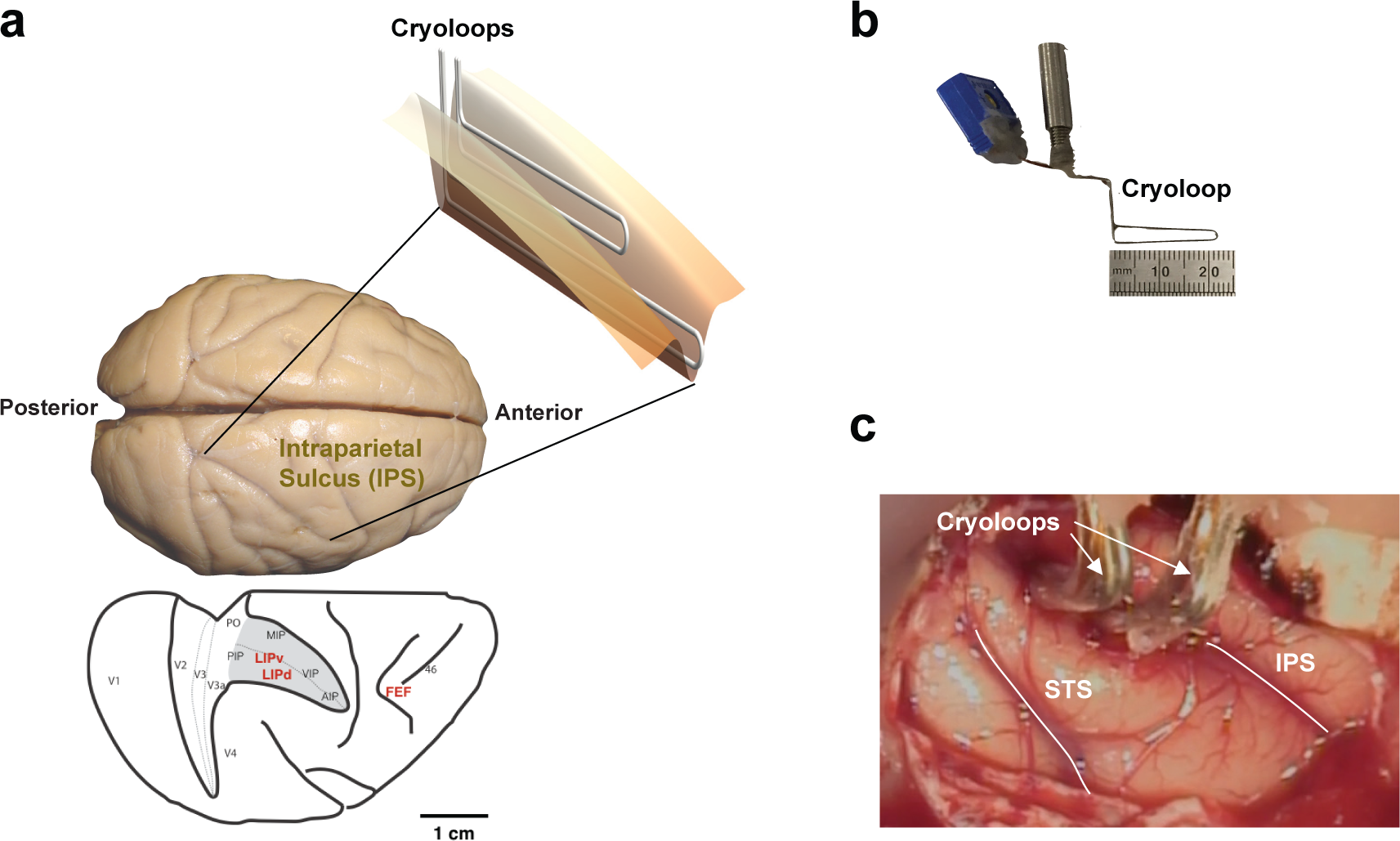
Implantation of cryoloops within the intraparietal sulcus (IPS). **a**. Schematic depiction of unilateral placement of two cryoloops within the IPS. Bottom outline of the brain shows the IPS opened, and the adjacent opened lunate sulcus. Shaded region shows the location of the cryloop implant amidst the surrounding areas of PPC. **b**. A single cryoloop custom-made to fit the full extent of the ventral half of the IPS. **c.** Intra-operative photograph of cryoloop implants in monkey J. Image shows both ventral and dorsal loops situated within the IPS, and the loops emerging from the sulcus, as well as the nearby superior temporal sulcus (STS). PO, posterior occipital area; PIP, posterior intraparietal area; MIP, medial intraparietal area; VIP, ventral intraparietal area; AIP, anterior intraparietal area; LIPv, ventral aspect of lateral intraparietal area; LIPd, dorsal aspect of lateral intraparietal area.

**Extended Data Figure 2.**
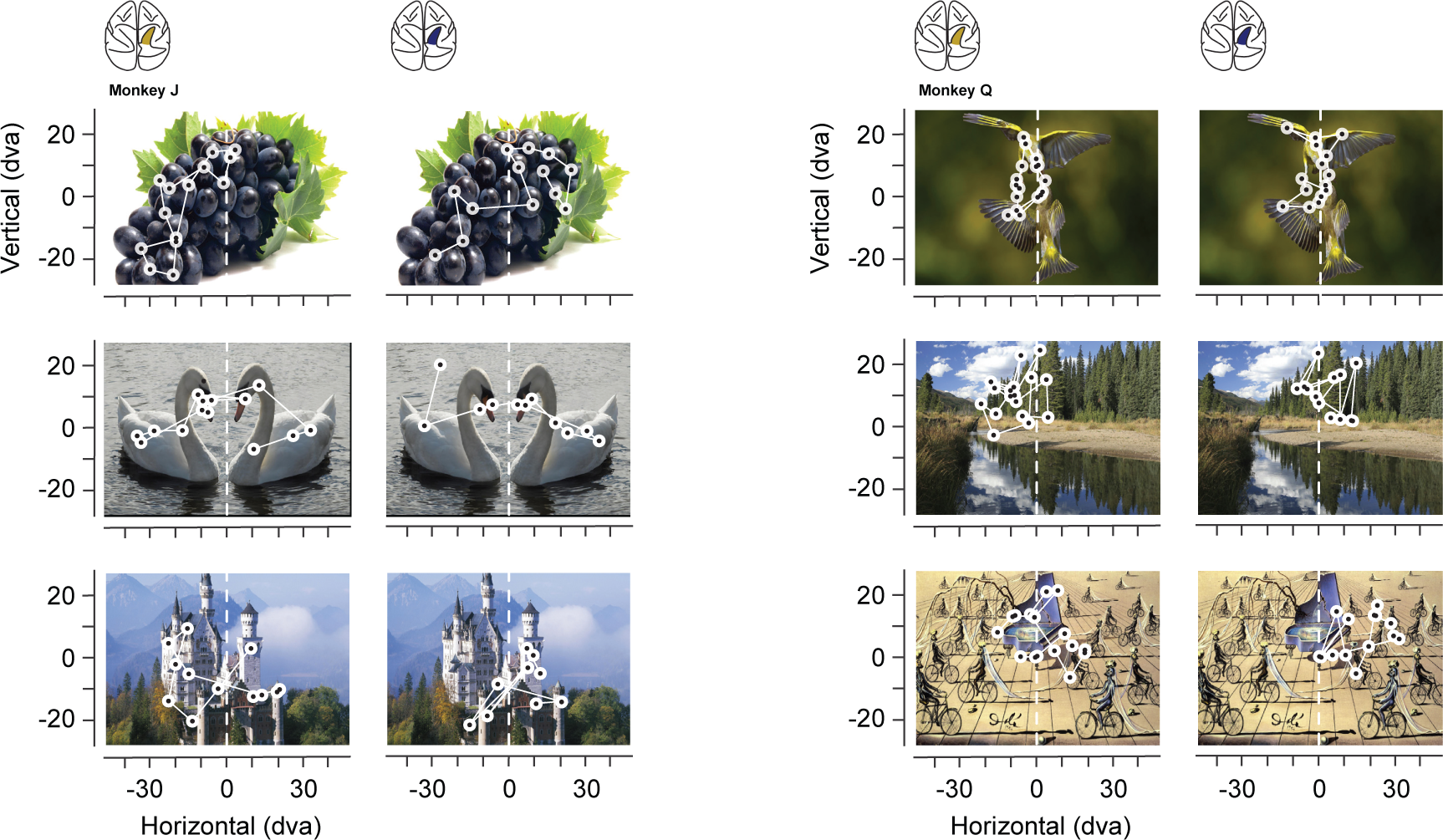
Free-viewing of images before and during parietal inactivation. Examples of images freely viewed by the two monkeys. Circles indicate regions of fixation and lines indicate saccades. For each monkey, control trials are shown on the left, and inactivation trials are shown on the right.

**Extended Data Figure 3.**
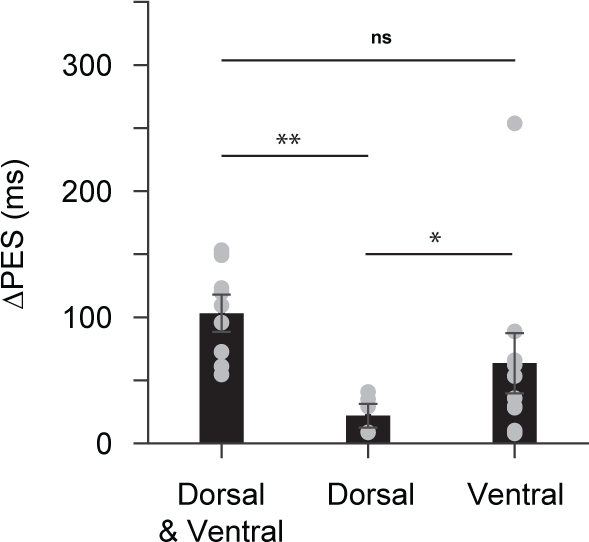
Behavioral effects of dorsal and ventral PPC inactivation. Comparisons of the shifts in double-target task effects during dorsal and ventral PPC inactivation in monkey Q. Bars show mean changes in PES values (inactivation – control) for sessions in which both dorsal and ventral cryoloops were cooled, or either the dorsal or ventral loops were cooled independently. Positive ΔPES values indicate a bias toward ipsilateral targets. Ventral cooling yielded effects comparable to cooling both dorsal and ventral loops and were more effective than dorsal cooling. Permutation was used for significance testing. Circles indicate data points from individual sessions. Error bars denote ±SEM; ns, not significant; *, P < 0.05; **, P < 0.01.

**Extended Data Figure 4.**
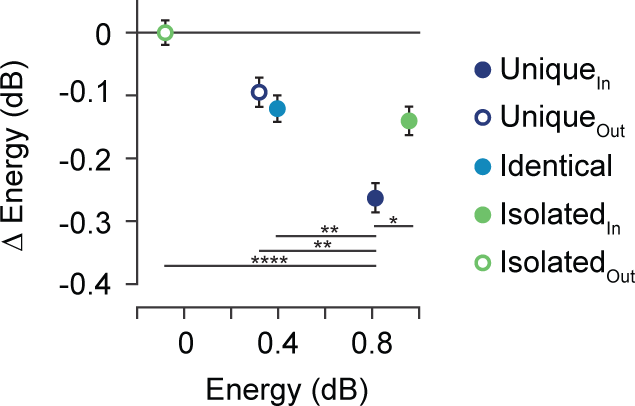
Changes in high-gamma band energy during PPC inactivation across different stimulus conditions. Points plot the mean high-gamma band LFP energy measured after the onset (0-500ms) of different visual stimuli. The reduction in high-gamma responses to unique stimuli in the URF was significantly larger than that of all other stimulus conditions (Unique_out_: p = 1.1 × 10^−3^; Isolated_In_: p = 0.027; Isolated_Out_: p < 10^−5^; Identical: p=1.6× 10^−3^). Paired permutation was used for significance testing. Error bars denote ±SEM; *, P < 0.05; **, P < 0.01; ****, P < 10^−4^.

**Extended Data Figure 5.**
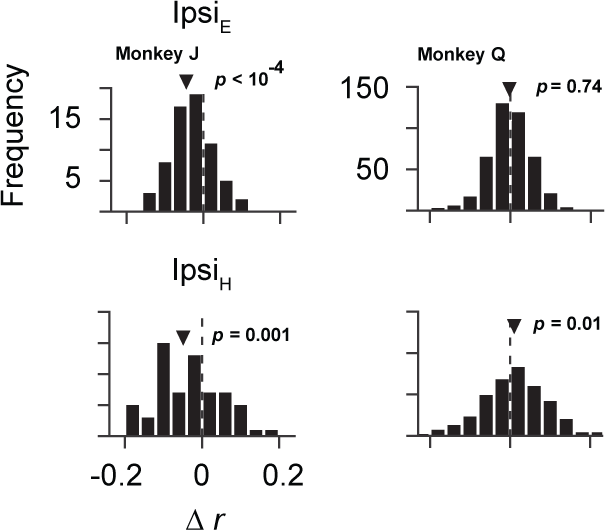
Changes in salience-guided fixations within the ipsilateral hemifield during PPC inactivation. **a**. Distribution of changes in fixation-salience map correlation coefficients (*r*_inactivation_ – *r*_control_) from ipsilateral fixations defined in eye-centered (Ipsi_E_) (top) or in head-centered (Ipsi_H_)(bottom) coordinates. Salience was computed using the GBVS model

**Extended Data Figure 6.**
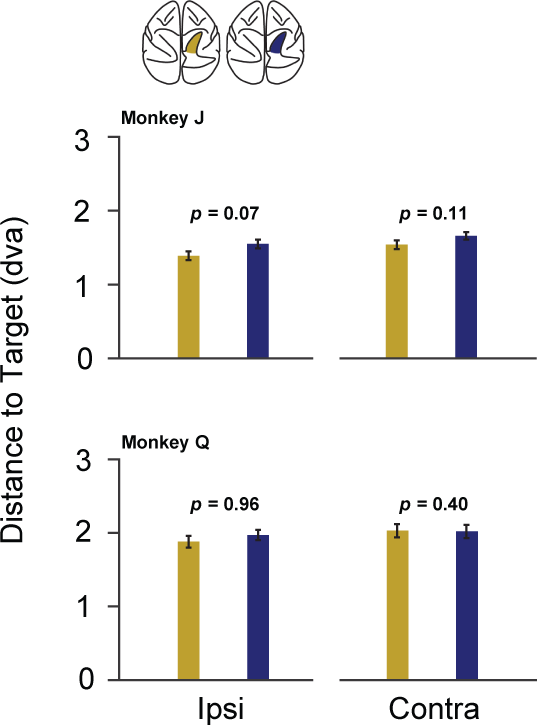
Effect of PPC inactivation on the accuracy of saccades to single targets. Bar plots show the mean saccadic error (distance to target) during control and inactivation trials for both monkeys. PPC inactivation did not significantly alter saccadic error in either hemifield of the two monkeys. Target eccentricity was 10 dva for monkey J and 15 dva for monkey Q. Error bars denote ±SEM.

**Extended Data Table 1.** Comparison of LFP energy between visual stimulus conditions for different frequency bands during control trials. LFP data were obtained after the onset [0-500) ms of different visual stimuli (n = 192 recordings). Paired t-tests were used to test significance.

| Frequency band | *M* (dB) | *SE* | *t*(191) | *P* |
| --- | --- | --- | --- | --- |
| **Isolated<sub>In</sub> vs Isolated<sub>Out</sub>** |  |  |  |  |
| Alpha | 1.436 | 0.161 | 8.903 | < 10<sup>-15</sup> |
| Beta | 0.034 | 0.090 | 0.375 | 0.708 |
| Gamma | 0.244 | 0.070 | 3.507 | < 10<sup>-3</sup> |
| High gamma | 1.052 | 0.058 | 18.065 | < 10<sup>-42</sup> |
| **Unique<sub>In</sub> vs Unique<sub>Out</sub>** |  |  |  |  |
| Alpha | 0.107 | 0.134 | 0.799 | 0.425 |
| Beta | -0.314 | 0.087 | -3.607 | < 10<sup>-3</sup> |
| Gamma | 0.136 | 0.067 | 2.038 | 0.043 |
| High gamma | 0.500 | 0.046 | 10.809 | < 10<sup>-20</sup> |
| **Unique<sub>In</sub> vs Identical** |  |  |  |  |
| Alpha | -0.113 | 0.132 | -0.860 | 0.391 |
| Beta | -0.540 | 0.102 | -5.308 | < 10<sup>-6</sup> |
| Gamma | 0.125 | 0.064 | 1.963 | 0.051 |
| High gamma | 0.469 | 0.042 | 11.075 | < 10<sup>-21</sup> |

**Extended Data Table 2.** Comparison of the reduction in visually driven activity across stimulus conditions during PPC inactivation. ANCOVA analysis examining the relationship between the change in visual activity during PPC inactivation (inactivation – control) and control visual activity across different visual stimulus conditions for all neuronal recordings (n = 352). Only the slope for the Unique-In condition differed significantly from the overall average.

|  | Intercept (spikes/s) | *P* | Slope | *P* |
| --- | --- | --- | --- | --- |
| **Average** | 0.88 | 0.01 | -0.08 | $< 10^{-35}$ |
| **Unique<sub>In</sub>** | 1.10 | 0.75 | -0.13 | $< 10^{-4}$ |
| **Unique<sub>Out</sub>** | 0.46 | 0.54 | -0.06 | 0.14 |
| **Identical** | 1.42 | 0.43 | -0.07 | 0.34 |
| **Isolated<sub>In</sub>** | -0.19 | 0.14 | -0.10 | 0.29 |
| **Isolated<sub>Out</sub>** | 1.60 | 0.28 | -0.06 | 0.06 |

## References

1. Moore, T. & Zirnsak, M. Neural Mechanisms of Selective Visual Attention. Annu. Rev. Psychol. 68, 47–72 (2017).

2. Bichot, N. P., Heard, M. T., DeGennaro, E. M. & Desimone, R. A Source for Feature-Based Attention in the Prefrontal Cortex. Neuron 88, 832–844 (2015).

3. Buschman, T. J. & Miller, E. K. Top-down versus bottom-up control of attention in the prefrontal and posterior parietal cortices. Science 315, 1860–1862 (2007).

4. Kastner, S., Pinsk, M. A., De Weerd, P., Desimone, R. & Ungerleider, L. G. Increased activity in human visual cortex during directed attention in the absence of visual stimulation. Neuron 22, 751–761 (1999).

5. Moore, T. & Fallah, M. Control of eye movements and spatial attention. Proc. Natl. Acad. Sci. 98, 1273–1276 (2001).

6. Ignashchenkova, A., Dicke, P. W., Haarmeier, T. & Thier, P. Neuron-specific contribution of the superior colliculus to overt and covert shifts of attention. Nat. Neurosci. 7, 56–64 (2004).

7. Krauzlis, R. J., Lovejoy, L. P. & Zénon, A. Superior colliculus and visual spatial attention. Annu. Rev. Neurosci. 36, 165–182 (2013).

8. Saalmann, Y. B., Pinsk, M. A., Wang, L., Li, X. & Kastner, S. The Pulvinar Regulates Information Transmission Between Cortical Areas Based on Attention Demands. Science 337, 753–756 (2012).

9. Zhou, H., Schafer, R. J. & Desimone, R. Pulvinar-Cortex Interactions in Vision and Attention. Neuron 89, 209–220 (2016).

10. Constantinidis, C. & Steinmetz, M. A. Neuronal responses in area 7a to multiple-stimulus displays: I. neurons encode the location of the salient stimulus. Cereb. Cortex N. Y. N 1991 11, 581–591 (2001).

11. Ipata, A. E., Gee, A. L., Gottlieb, J., Bisley, J. W. & Goldberg, M. E. LIP responses to a popout stimulus are reduced if it is overtly ignored. Nat. Neurosci. 9, 1071–1076 (2006).

12. Thompson, K. G. & Bichot, N. P. A visual salience map in the primate frontal eye field. Prog. Brain Res. 147, 251–262 (2005).

13. Allman, J., Miezin, F. & McGuinness, E. Stimulus specific responses from beyond the classical receptive field: neurophysiological mechanisms for local-global comparisons in visual neurons. Annu. Rev. Neurosci. 8, 407–430 (1985).

14. Burrows, B. E. & Moore, T. Influence and limitations of popout in the selection of salient visual stimuli by area V4 neurons. J. Neurosci. 29, 15169–15177 (2009).

15. Hegdé, J. & Felleman, D. J. How Selective Are V1 Cells for Pop-Out Stimuli? J. Neurosci. 23, 9968–9980 (2003).

16. Knierim, J. J. & van Essen, D. C. Neuronal responses to static texture patterns in area V1 of the alert macaque monkey. J. Neurophysiol. 67, 961–980 (1992).

17. Motter, B. C. Neural correlates of feature selective memory and pop-out in extrastriate area V4. J. Neurosci. Off. J. Soc. Neurosci. 14, 2190–2199 (1994).

18. Reynolds, J. H. & Desimone, R. Interacting roles of attention and visual salience in V4. Neuron 37, 853–863 (2003).

19. Hupé, J. M. et al. Cortical feedback improves discrimination between figure and background by V1, V2 and V3 neurons. Nature 394, 784–787 (1998).

20. Lomber, S. G., Payne, B. R. & Horel, J. A. The cryoloop: an adaptable reversible cooling deactivation method for behavioral or electrophysiological assessment of neural function. J. Neurosci. Methods 86, 179–194 (1999).

21. Ponce, C. R., Lomber, S. G. & Born, R. T. Integrating motion and depth via parallel pathways. Nat. Neurosci. 11, 216–223 (2008).

22. Smolyanskaya, A., Haefner, R. M., Lomber, S. G. & Born, R. T. A Modality-Specific Feedforward Component of Choice-Related Activity in MT. Neuron 87, 208–219 (2015).

23. Lynch, J. C. & McLaren, J. W. Deficits of visual attention and saccadic eye movements after lesions of parietooccipital cortex in monkeys. J. Neurophysiol. 61, 74–90 (1989).

24. Wardak, C., Olivier, E. & Duhamel, J.-R. Saccadic target selection deficits after lateral intraparietal area inactivation in monkeys. J. Neurosci. 22, 9877–9884 (2002).

25. Schiller, P. H. & Tehovnik, E. J. Cortical inhibitory circuits in eye-movement generation. Eur. J. Neurosci. 18, 3127–3133 (2003).

26. Soltani, A., Noudoost, B. & Moore, T. Dissociable dopaminergic control of saccadic target selection and its implications for reward modulation. Proc. Natl. Acad. Sci. 110, 3579–3584 (2013).

27. Liu, Y., Yttri, E. A. & Snyder, L. H. Intention and attention: different functional roles for LIPd and LIPv. Nat. Neurosci. 13, 495–500 (2010).

28. Vallar, G. Spatial hemineglect in humans. Trends Cogn. Sci. 2, 87–97 (1998).

29. Schall, J. D., Morel, A., King, D. J. & Bullier, J. Topography of visual cortex connections with frontal eye field in macaque: convergence and segregation of processing streams. J. Neurosci. 15, 4464–4487 (1995).

30. Lewis, J. W. & Van Essen, D. C. Corticocortical connections of visual, sensorimotor, and multimodal processing areas in the parietal lobe of the macaque monkey. J. Comp. Neurol. 428, 112–137 (2000).

31. Bichot, N. P., Schall, J. D. & Thompson, K. G. Visual feature selectivity in frontal eye fields induced by experience in mature macaques. Nature 381, 697–699 (1996).

32. Mohler, C. W., Goldberg, M. E. & Wurtz, R. H. Visual receptive fields of frontal eye field neurons. Brain Res. 61, 385–389 (1973).

33. Chen, X., Zirnsak, M. & Moore, T. Dissonant Representations of Visual Space in Prefrontal Cortex during Eye Movements. Cell Rep. 22, 2039–2052 (2018).

34. Schall, J. D. On the role of frontal eye field in guiding attention and saccades. Vision Res. 44, 1453–1467 (2004).

35. Schiller, P. H., True, S. D. & Conway, J. L. Effects of frontal eye field and superior colliculus ablations on eye movements. Science 206, 590–592 (1979).

36. Koch, C. & Ullman, S. Shifts in selective visual attention: towards the underlying neural circuitry. Hum. Neurobiol. 4, 219–227 (1985).

37. Harel, J., Koch, C. & Perona, P. Graph-based visual saliency. in Advances in neural information processing systems 545–552 (2007).

38. Itti, L., Koch, C. & Niebur, E. A model of saliency-based visual attention for rapid scene analysis. IEEE Trans. Pattern Anal. Mach. Intell. 1254–1259 (1998).

39. Wang, J., Borji, A., Kuo, C.-J. & Itti, L. Learning a Combined Model of Visual Saliency for Fixation Prediction. IEEE Trans. Image Process. 25, 1566–1579 (2016).

40. Borji, A., Sihite, D. N. & Itti, L. Quantitative Analysis of Human-Model Agreement in Visual Saliency Modeling: A Comparative Study. IEEE Trans. Image Process. 22, 55–69 (2013).

41. Gottlieb, J. P., Kusunoki, M. & Goldberg, M. E. The representation of visual salience in monkey parietal cortex. Nature 391, 481–484 (1998).

42. Goldring, A. & Krubitzer, L. Evolution of parietal cortex in mammals: From manipulation to tool use. in The Evolution of Nervous Systems 3, 259–296 (Elsevier, 2017).

43. Soltani, A. & Koch, C. Visual saliency computations: mechanisms, constraints, and the effect of feedback. J. Neurosci. Off. J. Soc. Neurosci. 30, 12831–12843 (2010).

44. De Weerd, P., Desimone, R. & Ungerleider, L. G. Generalized deficits in visual selective attention after V4 and TEO lesions in macaques. Eur. J. Neurosci. 18, 1671–1691 (2003).

45. Gregoriou, G. G., Rossi, A. F., Ungerleider, L. G. & Desimone, R. Lesions of prefrontal cortex reduce attentional modulation of neuronal responses and synchrony in V4. Nat. Neurosci. 17, 1003 (2014).

46. Newsome, W. T. & Paré, E. B. A selective impairment of motion perception following lesions of the middle temporal visual area (MT). J. Neurosci. Off. J. Soc. Neurosci. 8, 2201–2211 (1988).

47. White, B. J., Kan, J. Y., Levy, R., Itti, L. & Munoz, D. P. Superior colliculus encodes visual saliency before the primary visual cortex. Proc. Natl. Acad. Sci. U. S. A. 114, 9451–9456 (2017).

48. Mysore, S. P. & Knudsen, E. I. A shared inhibitory circuit for both exogenous and endogenous control of stimulus selection. Nat. Neurosci. 16, 473 (2013).

49. Lomber, S. G. & Payne, B. R. Translaminar differentiation of visually guided behaviors revealed by restricted cerebral cooling deactivation. Cereb. Cortex 10, 1066–1077 (2000).

50. Lomber, S. G., Cornwell, P., Sun, J. S., MacNeil, M. A. & Payne, B. R. Reversible inactivation of visual processing operations in middle suprasylvian cortex of the behaving cat. Proc. Natl. Acad. Sci. U. S. A. 91, 2999–3003 (1994).

51. Lomber, S. G., Payne, B. R., Cornwell, P. & Long, K. D. Perceptual and cognitive visual functions of parietal and temporal cortices in the cat. Cereb. Cortex 6, 673–695 (1996).

52. Killian, N. J., Jutras, M. J. & Buffalo, E. A. A map of visual space in the primate entorhinal cortex. Nature 491, 761–764 (2012).

53. Noudoost, B. & Moore, T. Control of visual cortical signals by prefrontal dopamine. Nature 474, 372 (2011).

54. Chen, X. & Stuphorn, V. Inactivation of Medial Frontal Cortex Changes Risk Preference. Curr. Biol. (2018).

55. Chang, C.-C. & Lin, C.-J. LIBSVM: A library for support vector machines. ACM Trans. Intell. Syst. Technol. TIST 2, 27 (2011).

56. Chandran K S, S., Mishra, A., Shirhatti, V. & Ray, S. Comparison of Matching Pursuit Algorithm with Other Signal Processing Techniques for Computation of the Time-Frequency Power Spectrum of Brain Signals. J. Neurosci. Off. J. Soc. Neurosci. 36, 3399–3408 (2016).

57. Chen, X., Scangos, K. W. & Stuphorn, V. Supplementary motor area exerts proactive and reactive control of arm movements. J. Neurosci. Off. J. Soc. Neurosci. 30, 14657–14675 (2010).

